# Collagen Density Modulates the Immunosuppressive Functions of Tumor-Associated Macrophages

**DOI:** 10.1101/513986

**Authors:** Anne Mette H. Larsen, Dorota E. Kuczek, Adrija Kalvisa, Majken S. Siersbæk, Marie-Louise Thorseth, Astrid Zedlitz Johansen, Marco Carretta, Lars Grøntved, Ole Vang, Daniel H. Madsen

## Abstract

Tumor-associated macrophages (TAMs) support tumor growth by suppressing the activity of tumor infiltrating T cells. Consistently, the number of TAMs has been correlated with a poor prognosis of cancer. The immunosuppressive TAMs are also considered a major limitation for the efficacy of cancer immunotherapy. However, the molecular reason behind the acquisition of an immunosuppressive TAM phenotype is still not completely understood. During solid tumor growth, the extracellular matrix (ECM) is degraded and substituted with a tumor specific collagen-rich ECM. The collagen density of this tumor ECM has been associated with a poor prognosis of several cancers, but the underlying reason for this correlation is not well understood. Here, we have investigated whether the collagen density could modulate the immunosuppressive activity of TAMs and thereby promote tumor progression.

In this study, the macrophage cell line RAW 264.7 was 3D cultured in collagen matrices of low- and high collagen densities mimicking healthy and tumor tissue, respectively. The effects of collagen density on macrophage phenotype and function were investigated by confocal microscopy, flow cytometry, RNA sequencing, qRT-PCR, and ELISA analysis. To investigate the effect of collagen density on the immune modulatory activity of macrophages, co-culture assays with primary T cells to assess T cell chemotaxis and proliferation were conducted. Lastly, the effects of collagen density on primary cells were investigated using murine bone-marrow derived macrophages (BMDMs) and TAMs isolated from murine 4T1 breast tumors.

Collagen density did not affect the proliferation, viability or morphology of macrophages. However, whole-transcriptome analysis revealed a striking response to the surrounding collagen density including the differential regulation of many immune regulatory genes and genes encoding chemokines. The transcriptional changes in RAW 264.7 macrophages were shown to be similar in murine BMDMs and TAMs. Strikingly, the collagen density-induced changes in the gene expression profile had functional consequences for the macrophages. Specifically, macrophages cultured in high density collagen were less efficient at attracting cytotoxic T cells and also capable of inhibiting T cell proliferation to a greater extent than macrophages cultured in low density collagen.

Our study demonstrates that a high collagen density can instruct TAMs to acquire an immunosuppressive phenotype. This could be one of the mechanisms decreasing the efficacy of immunotherapy and linking increased collagen density to poor patient prognosis.

## Introduction

Immunotherapy is a collection of several promising ways of treating cancer, which all use the host’s own immune response to fight the disease. The various kinds of immunotherapies use different strategies to induce T cells to kill the cancer cells^1,2^. Checkpoint-inhibitor therapy (anti-CTLA4 and anti-PD-1/PD-L1) and adoptive T cell transfer are two of the most successful types of immunotherapies^1,2^. They both exploit the ability of endogenous T cells to recognize and kill cancer cells. Unfortunately, not all patients respond to immunotherapy and some patients only display a partial response followed by relapse^3^. A likely explanation is the existence of a highly immunosuppressive tumor microenvironment (TME), which inhibits the activity of tumor-infiltrating T cells and thereby reduces the therapeutic efficacy^4,5^. The mechanisms underlying the generation of an immunosuppressive TME are still only partially understood.

During solid tumor growth, the surrounding extracellular matrix (ECM) is degraded and substituted with a tumor specific ECM. This ECM often contains high amounts of collagen type I, is more linearized and of increased stiffness^6,7^. The density and structure of the tumor ECM have been linked to a poor prognosis of several cancers such as breast cancer^8^, pancreatic cancer^9^, gastric cancer^10^, and oral squamous cell carcinomas^11^, but the reason for this correlation is still not clear. It has been demonstrated *in vitro* that the density and stiffness of the ECM can affect various cell types, including mesenchymal stem cells^12^, fibroblasts^13,14^, T cells^15^, and cancer cells^7–16,17^. This includes the differentiation of mesenchymal stem cells towards a neurogenic or osteogenic lineage when cultured on soft or stiff matrices, respectively^12^, and the stimulation of malignant transformation of two breast epithelial cell lines when cultured on stiff matrices^16^. However, still very little is known about the effect of ECM on the immune cells. In this regard, the tumor-associated macrophages (TAMs) are of particular interest due to their pro-tumorigenic activity.

Macrophages are known to be plastic cells that, depending on the environment, can acquire different phenotypes, becoming either pro-or anti-inflammatory^18^. The pro-inflammatory macrophages are also referred to as classically activated-or M1-polarized macrophages^19^. They are characterized by the expression of numerous pro-inflammatory cytokines such as *IL-12, iNOS* and *IL-6*^19^. Anti-inflammatory macrophages are also referred to as alternatively activated macrophages or M2-polarized macrophages. They are characterized by the expression of several immunosuppressive factors such as *IL10, CCL17* and CCL22^19^. In addition to being anti-inflammatory, they can promote tissue remodeling, angiogenesis, and wound healing by digestion of the ECM^20–22^.

TAMs often acquire an anti-inflammatory M2-like phenotype and are known to support tumor growth through various mechanisms including the suppression of tumor-infiltrating T cells^23–25^. This immunosuppressive activity of TAMs is considered a major limitation for immunotherapy efficacy, and consistently, the number of TAMs correlates with a poor prognosis in many types of cancer^26–28^. Therapeutic approaches aiming at re-programming TAMs toward a more pro-inflammatory phenotype have already been tested in mice^29,30^. These have shown promising results in various cancers, and have stimulated the initiation of clinical trials^31^. However, the molecular reason why TAMs gain an immunosuppressive phenotype in the TME is still not completely understood.

A previous study has shown that a larger percentage of human monocytes differentiate into macrophages when cultured on collagen type I coated dishes compared to when cultured on uncoated dishes^32^. Another study has shown that alveolar macrophages cultured in the presence of collagen type I monomers acquire a more anti-inflammatory phenotype^33^. However, in this study macrophages were subjected to soluble collagen monomers and not native 3D collagen matrices. Recently, it was demonstrated that implantation of a collagen gel into an injured muscle of a mouse, stimulated macrophages to become more immunosuppressive^34^. The studies illustrate that collagen can affect macrophages both *in vitro* and *in vivo* and suggest that collagen could influence the immunosuppressive properties of TAMs in the TME.

In this study, we have used 3D culture assays to investigate if collagen density can induce immunosuppressive activity of macrophages.

## Materials and Methods

### RAW 264.7 culture in 3D collagen matrices

The murine macrophage cell line RAW 264.7 was cultured in RPMI supplemented with 10% FBS and 1% P/S for no more than 20 passages and split one day prior to 3D culture in collagen matrices. 3D type I collagen matrices were prepared based on the protocol by Artym and Matsumoto^35^. Briefly, collagen matrices were generated by mixing rat tail collagen type I (Corning), 0.02N acetic acid, 10x DMEM with phenol red (Sigma Aldrich) and 10x reconstitution buffer (0.2M Hepes (Gibco) and 0. 262M NaHCO3). To neutralize the pH, 2N NaOH was added to the reconstitution buffer prior to use. First, 350 μl collagen solution of either low-(1 mg/ml) or high (4 mg/ml) collagen density was prepared and added to wells of non-tissue-culture treated 24-well plates (Corning). The collagen solution was allowed to polymerize for at least 30 min. at 37°C, 5% CO_2_. Next, 350 μl collagen solution containing 60,000 RAW 264.7 macrophages was added onto the first collagen gel of matching concentration. The gel was allowed to polymerize for 2-3h after which 600 μl prewarmed RPMI supplemented with 10% FBS and 1% P/S was added on top of the collagen gel. To remove salts and potentially unpolymerized collagen present, the media on all samples was changed twice after 24h of culture. For conventional 2D culture, 60,000 cells were seeded per well of tissue-culture treated 6-well plates.

### Confocal microscopy

3D collagen matrices of 1 mg/ml or 4 mg/ml collagen were made as previously described but with additionally 20% acid-extracted rat tail tendon type I collagen labelled with Alexa Fluor 647 incorporated. The Alexa Fluor labelling was done as previously described^21^. A total volume of 200 μl collagen solution containing 60,000 RAW 264.7 macrophages was added per well of a glass bottom 24-well, No. 0 Coverslip, 13 mm Glass Diameter plate (MatTek Corporation) and allowed to polymerize for 45 min. at 37°C, 5% CO_2_. Thereafter, 500 μl of RPMI supplemented with 10% FBS and 1% P/S was added to the matrices. Prior to microscopy, samples were stained as described by Artym and Matsumoto^35^. Briefly, samples were fixed with 4% formaldehyde, 5% sucrose for 1h and permeabilized with 0.5% Triton-X for 10 min. Next, F-actin and nuclei were stained with 1:40 diluted Alexa Fluor^®^ 488 Phalloidine (Thermo Fisher Scientific) and 1:1000 diluted DAPI (Sigma Aldrich) for 1h and 20 min., respectively. Confocal imaging was performed using an IX83 Olympus confocal microscope equipped with a scanning head (FluoView 1200; Olympus). All images were acquired using an Olympus UPlanSApo 60x/1.35na oil objective.

### Proliferation and viability of RAW 264.7 macrophages

The proliferation of RAW 264.7 macrophages was determined using an APC BrdU Flow Kit (BD Biosciences) according to the manufacturer’s protocol. Briefly, RAW 264.7 macrophages were cultured for three days in 3D collagen matrices of 1 mg/ml or 4 mg/ml collagen as previously described or on 2D tissue-culture treated 6-well plates. To label the cells, all samples were pulsed with a final concentration of 10 μM BrdU for 1h. Following labelling, culture media was aspirated from both 2D and 3D samples, and all samples were treated with 500 μl 3 mg/ml collagenase type I (Worthington) for 45-60 min. at 37°C, 5% CO_2_ to dissolve the collagen matrices. Cell solutions were collected and washed once with DPBS (Lonza) before staining with Zombie Aqua Fixable Viability Dye (BioLegend). Next, cells were fixed, permeabilized and stained with APC anti-BrdU antibody according to the manufacturer’s protocol (BD Biosciences). Cells were resuspended in FACS buffer (DPBS with 5 mM EDTA and 0.5% BSA) and run on a BD^TM^ LSR II flow cytometer. Data were analyzed using FACS Diva^®^ 8.0.2 software.

### RNA sequencing

RAW 264.7 macrophages were isolated from 3D collagen matrices of 1 mg/ml or 4 mg/ml collagen and from 2D tissue-culture treated 6-well plates. RNA was extracted using the RNeasy Mini Kit (Qiagen). The quantity and purity of isolated RNA were assessed using an Agilent 2100 BioAnalyzer (Agilent Genomics). Total RNA (1000 ng) with RIN values > 9.9 were used for RNA sequencing. Libraries were synthesized from PolydT enriched mRNA using random priming according to the NEBNext RNA Library Preparation Kit for Illumina. Library quality was assessed using Fragment Analyzer (AATI), followed by library quantification (KAPA). Sequencing was done on a HiSeq1500 platform (Illumina) with a read length of 50 bp. Sequenced reads were aligned to mus musculus mm9 using STAR^36^. Uniquely aligned reads were quantified at exons of annotated genes and normalized to sequence depth and gene length using HOMER^37^. The number of reads per kilobase per million mapped (RPKM) for all RefSeq annotated genes can be found in Table S1. The analysis of differential expression was performed using the DESeq2 package^38^ in R. Principal component analysis was performed using R (prcomp package). Gene ontology analysis were performed by GOseq^39^ in R, using only data with FDR < 0.05 and Log2 fold change > 0.585. Heatmaps were generated from z-score normalized RPKM values using R (pheatmap package) on selected sets of genes. MA-plots and Volcano plots were generated using R.

### RNA extraction, cDNA synthesis and quantitative real-time-PCR (qRT-PCR)

RNA was isolated from RAW 264.7 macrophages cultured in 3D collagen matrices of 1 mg/ml or 4 mg/ml collagen as previously described. The quantity and purity of isolated RNA were assessed using an Agilent 2100 BioAnalyzer (Agilent Genomics). Samples with RIN values > 9.4 were used for further analysis. cDNA was generated using iScript™ cDNA Synthesis Kit (Bio Rad). As a control, a noRT sample was included during the cDNA synthesis, containing all components but the reverse transcriptase. The qRT-PCR reaction was made using the Brilliant III Ultra-Fast SYBR^®^ Green QPCR Master Mix (Agilent Technologies). Briefly, the reaction included an initial activation step at 95°C for 3 min., then 40 cycles of denaturation at 95°C for 5 sec., and annealing/extension at 60°C for 20 sec., and lastly a high resolution melting curve analysis of 65–95°C with 0.5°C increment, 5 sec. per step. Equal amounts of cDNA were added to all samples. As a negative control, a sample containing all qRT-PCR components except the cDNA template was included in the analysis. All samples were run on an AriaMX Real Time PCR system (G8830A). All measurements were based on quadruplicates of each cell culture condition and normalized to the internal reference gene, *Actb.* The comparative cycle threshold (ΔΔCT) method was used to calculate the relative fold changes. Primers were designed using the Primer-BLAST tool (NCBI, NIH). Only primers spanning exon-exon junction, and with a maximum product length of 250 bp was used. Prior to use, the primer efficiency for all primer sets were measured and found to be between 86% and 111%. Primer sequences are listed in Table S2.

### CCL3 and PGE2 ELISA

ELISAs were performed using Mouse MIP-1 alpha (CCL3) ELISA Ready-SET-Go ELISA kit (eBi-oscience™) and Prostaglandin E2 ELISA Kit (Abcam) according to the manufacturer’s instructions. Analysis was done on conditioned media from RAW 264.7 macrophages cultured in 3D collagen matrices of 1 mg/ml or 4 mg/ml collagen. To evaluate the amount of collagen-bound protein, 3D collagen samples were treated with collagenase type I as previously described. All samples were centrifuged for 5 min. at 300 x g and stored at −20°C prior to use. All measurements were based on quadruplicates of each cell culture condition.

### T cell isolation

T cells were isolated from spleens of BALB/c mice using CD3ε MicroBead Kit, mouse (Miltenyi Biotec). Briefly, single cell suspensions were generated by forcing freshly isolated spleens through 70 μm cell strainers using the piston of a syringe. Red blood cells were lysed using Red Blood Cell Lysis buffer (Qiagen) for 1 min. at room temperature. The T cell isolation was done according to the manufacturer’s protocol (Miltenyi Biotec), however prolonging the incubation with CD3ε-Biotin to 20 min. and anti-Biotin to 25 min. Following the isolation, small fractions of the CD3 negative and positive cells were stained with CD3-FITC (BioLegend) and Zombie Aqua Fixable Viability Dye (BioLegend) and run on a BD^TM^ LSR II flow cytometer to evaluate the purity. A purity >80% living CD3+ cells was considered satisfactory for use in subsequent experiments. A total of 4×10^6^ isolated T cells per well were rested O.N. in RPMI supplemented with 10% FBS, 1% P/S, 50 μM 2-mercaptoethanol, 0.1 mM non-essential amino acids, 1mM Sodium Pyruvate (From now on referred to at T cell media), and 3-5 μg/ml Concanavalin A (ConA) (Sigma Aldrich) in tissue-culture treated 24-well plates.

### T cell proliferation

For co-culture assays, splenocytes were isolated from BALB/c mice as previously described. RAW 264.7 macrophages were cultured in 3D collagen matrices of 1 mg/ml or 4 mg/ml collagen for one day prior to co-culture. On the day of co-culture, the RAW 264.7 macrophages were washed once with T cell media after which 600 μl T cell media supplemented with 3-5 μg/ml ConA (Sigma Aldrich) was added. A total of 400,000 freshly isolated splenocytes in T cell media supplemented with 3-5 μg/ml ConA (Sigma Aldrich) were added to 6.5 mm transwell inserts with 0.4 μm pore size polyester membranes (Corning) placed above the 3D collagen matrices, corresponding to a 3:1 ratio of T cells to RAW 264.7 macrophages. As controls, splenocytes were cultured alone in T cell media supplemented with 3-5 μg/ml ConA (Sigma Aldrich) or together with collagen matrices of 1 mg/ml or 4 mg/ml collagen without any embedded RAW 264.7 macrophages. Cells were co-cultured for three days. T cell proliferation was determined using an APC BrdU Flow Kit (BD Biosciences) according to the manufacturer’s protocol. Briefly, the splenocytes were pulsed with a final concentration of 10 μM BrdU for 1h and collected from the transwell inserts. Cells were washed once with DPBS (Lonza) before staining with Zombie Aqua Fixable Viability Dye, CD3-FITC, CD4-BV421, CD8-APC-Cy7, CD25-PerCP-Cy5.5, and PD1-PE (All BioLegend). Next, cells were fixed, permea-bilized and stained with APC anti-BrdU antibody according to the manufacturer’s protocol (BD Bioscience). Cells were resuspended in FACS buffer and run on a BD^TM^ LSR II or an ACEA NovoCyte Quanteon™ flow cytometer. Data were analyzed using FACS Diva^®^ 8.0.2 or NovoExpress^®^ software. Measurements were based on triplicates or quadruplicates of each cell culture condition.

### T cell migration

The migration of T cells towards conditioned media from RAW 264.7 macrophages was measured using O.N. rested primary murine T cell isolated as previously described. Conditioned media was harvested from RAW 264.7 macrophages grown for three days in collagen matrices of 1 mg/ml or 4 mg/ml collagen with 600 μl T cell media. A total of 300,000 T cells in 200 μl T cell media supplemented with 3-5 μg/ml ConA (Sigma Aldrich) were added to 6.5 mm transwell inserts with 5.0 μm pore size polycarbonate membranes (Corning). The T cells were placed above 500 μl conditioned media supplemented with 3-5 μg/ml ConA (Sigma Aldrich) in 24-well plates. Prior to use, conditioned media was centrifuged at 300 x g for 5 min. and ConA (Sigma Aldrich) was added to a final concentration of 3-5 μg/ml. As a control for maximal migration, 300,000 T cells were added directly to the wells of a 24-well plate. The T cells were allowed to migrate towards the conditioned media for 26-28h at 37°C, 5% CO_2_. Subsequently, all migrated cells were collected from the lower wells and stained with CD3-FITC, CD4-BV421, CD8-APC, CD25-PerCP-Cy5.5 and Zombie Aqua Fixable Viability Dye (All BioLegend). The wells were additionally washed once with FACS buffer to ensure the collection of all cells. To determine the number of migrated cells, all samples were resuspended in 200 μl FACS Buffer and run on a BD^TM^ LSR II Flow Cytometer for 2 min. on flow rate “high”. Specific migration was determined by normalizing the data to the maximum control.

### 3D culture of murine bone-marrow derived macrophages (BMDMs)

To investigate the effect of collagen density on BMDMs, femurs and tibias were isolated from BALB/c mice. Single cell suspensions were established by flushing the bones with ice cold DPBS (Lonza) using a syringe and passing the resulting cells through a 70 μm cell strainer. Red blood cells were removed by treating the cells with 5 ml Red Blood Cell Lysis buffer (Qiagen) for 4 min. Next, the cells were resuspended to a concentration of 400,000 cells/ml in DMEM supplemented with 20% FBS, 1% P/S and 20 ng/ml M-CSF (ReproTech). The cells were plated on non-tissue-culture treated plates for six days, replenishing the media on day three. On day six, M-CSF stimulated BMDMs were harvested by incubation in ice cold 4 mM EDTA in PBS for 10 min. while occasionally pipetting up and down. The purity of the cells was assessed by staining a small fraction with Zombie Aqua Fixable Viability Dye, F4/80-APC and CD11b-Pe-Cy7 (all Biolegend) and analyzing them on a BD^TM^ LSR II Flow Cytometer. A total of 600,000 M-CSF stimulated BMDMs were cultured in 3D matrices of 1 mg/ml or 4 mg/ml collagen with DMEM supplemented with 20% FBS, 1% P/S added on top. The cells were cultured for one day before isolating RNA (Qiagen) as previously described. A total amount of 400-1000 ng RNA was converted to cDNA using iScript™ cDNA Synthesis Kit (Bio Rad). qRT-PCR reactions were made for selected genes using the Brilliant III Ultra-Fast SYBR^®^ Green QPCR Master Mix (Agilent Technologies) as previously described.

### 3D culture of murine TAMs

To generate murine tumors, 500,000 4T1 cancer cells in 100 μl RPMI were subcutaneously injected into each side of the flanks of BALB/c mice. Tumors were allowed to grow to a maximum size of 864 mm^3^, after which the mice were euthanized. Tumors were isolated, dissected into smaller fragments and digested in RPMI supplemented with 2.1 mg/ml collagenase type I, 75 μg/ml DNase I, 5mM CaCl2 and 1% P/S at 4°C, O.N and then at 37°C for 30-60 min. Single cell suspensions were then established by resuspending the cells and passing them through a 70 μm cell strainer. Red blood cells were lysed by treating the cells with Red Blood Cell lysis buffer (Qiagen) for 5 min. Next, F4/80+ TAMs were isolated from the cell suspension using anti-F4/80 MicroBeads (Miltenyi Biotec) according to the manufacturer’s protocol. The purity of the isolated TAMs was assessed by staining a small fraction with Zombie Aqua Fixable Viability Dye, F4/80-APC and CD11b-Pe-Cy7 (all Biolegend) and analyzing them on a BD^TM^ LSR II flow cytometer. Following isolation, 600,000 F4/80+ TAMs were cultured in collagen matrices of 1 mg/ml or 4 mg/ml collagen densities with DMEM supplemented with 20% FBS and 1% P/S added on top. The cells were cultured for one day before isolating RNA (Qiagen) as previously described. A total amount of 500-750 ng RNA was converted into cDNA using iScript™ cDNA Synthesis Kit (Bio Rad). qRT-PCR reactions were made for selected genes using the Brilliant III Ultra-Fast SYBR^®^ Green QPCR Master Mix (Agilent Technologies) as previously described.

### Statistical analysis

All experiments were performed at least three times with at least three replicates per condition. All results are presented as the mean +/− standard deviation unless otherwise specified. For two-group comparisons of the means from independent experiments, paired two-tailed Student’s t-tests were performed unless otherwise specified. For multi-group comparisons, one-way analysis of variance (ANOVA) was used followed by un-paired two-tailed Student’s t-tests unless otherwise specified. All statistical analyses, except RNASeq data, were performed using GraphPad Prism for Windows, Version 7. A p-value<0.05 was considered statistically significant. Statistical analysis of RNASeq data was done using R.

## Results

### Macrophage viability and proliferation is unaffected by the surrounding collagen density

To investigate if 3D culture of macrophages in collagen matrices of different densities affects the viability and proliferation of macrophages, the murine macrophage cell line RAW 264.7 was embedded and cultured in collagen type I matrices of low (1 mg/ml) or high (4 mg/ml) collagen density or on regular tissue-culture treated plastic (2D culture). The selected collagen concentration of 1 mg/ml is representative of healthy normal tissue such as lung or mammary gland whereas 4 mg/ml collagen mimics the ECM density of solid tumors^16,40^. To ensure 3D cultured cells were not in contact with any plastic, a lower layer of polymerized collagen was generated before adding a second layer of collagen containing RAW 264.7 macrophages. Using confocal microscopy, we confirmed that the cells were completely surrounded by collagen fibers and located in different planar levels of the matrix (Figure 1A). Additionally, it was clearly seen that the structure and density of collagen were different when comparing low and high collagen density matrices (Figure 1B-C). However, when investigating the morphology of the macrophages grown in collagen matrices of low or high density no obvious differences were observed (Figure 1B-C). To evaluate if 3D culture affected the cellular viability, RAW 264.7 macrophages were cultured in collagen matrices of low- or high density, stained with a live/dead marker and analyzed by flow cytometry. 3D culture of RAW 264.7 macrophages in collagen matrices did not affect the viability of the cells compared to cells subjected to conventional 2D culture on a plastic surface (Figure 1D). Additionally, no difference in viability was observed between macrophages cultured in low- or high density collagen (Figure 1D). Next, the proliferation of the macrophages was evaluated using a BrdU-based flow cytometry assay. Neither culture of the macrophages in 3D compared to 2D nor culture of the macrophages in 3D collagen matrices of different densities affected the proliferation rate of the RAW 264.7 macrophages (Figure 1E).

**Figure 1.**
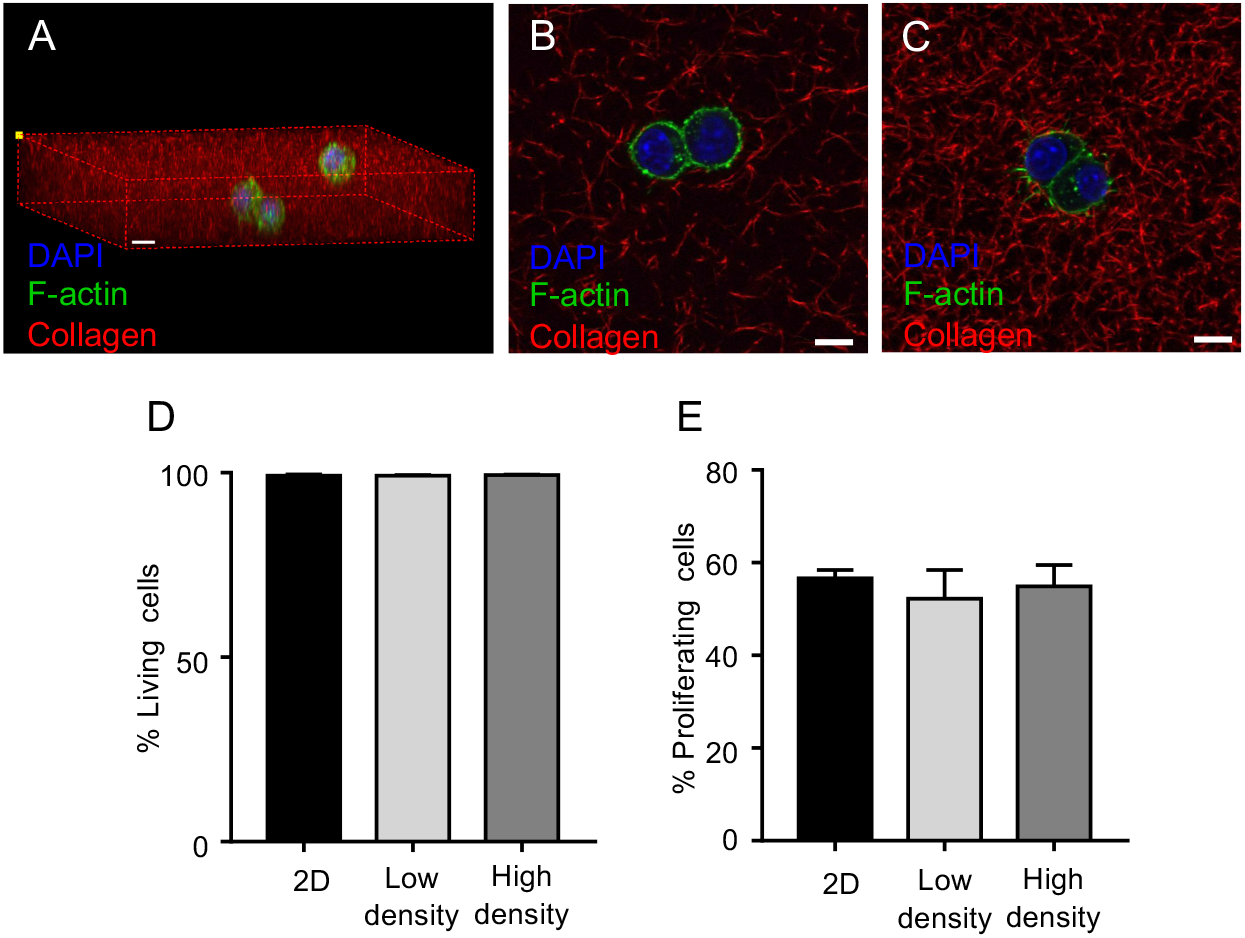
Collagen density does not affect viability and proliferation of RAW 264.7 macrophages. (A-C) Representative confocal microscopy images of RAW 264.7 macrophages grown in 3D collagen matrices of 1 mg/ml (Low density, A and B) or 4 mg/ml (High density, C) collagen. (A) 3D projection of collagen-embedded RAW 264.7 macrophages. (B and C) Single sections of RAW 264.7 macrophages embedded in collagen. (A-C) Scale bars = 10 μm. (D and E) RAW 264.7 macrophages were cultured for three days in collagen matrices of low- or high density or on tissue-culture treated plastic (2D) and analyzed by flow cytometry to assess viability (D) and proliferation (E). The results are based on three individual experiments with three samples per condition in each. Error bars = SEM.

### 3D culture changes the gene expression profile of macrophages

Although 3D culture in different collagen densities did not affect the morphology, viability, or proliferation of RAW 264.7 macrophages, we speculated that the transcriptional profile of the macrophages could still be affected. To investigate this possibility, RNA was extracted from RAW 264.7 macrophages cultured in collagen matrices of low- or high densities or on tissue-culture treated plastic and subjected to RNA sequencing. A principal component analysis of the resulting data showed that the transcriptional profile of macrophages cultured in 3D (low- or high density collagen) separated from the 2D cultured macrophages (Figure 2A). However, strikingly also that macrophages cultured in high density collagen clustered separately from macrophages cultured in low density collagen (Figure 2A). Additional analysis of the difference between 3D and 2D culture revealed that 1271 and 1490 genes (FDR<0.01, Log2 fold change > 0.585) were differentially regulated when comparing macrophages cultured on 2D tissue-culture treated plates with macrophages cultured in 3D low density collagen (Figure 2B, Figure S1A and Table S3) and 3D high density collagen (Figure S1B-S2A and Table S4), respectively. The full gene expression dataset is in Table S1. To investigate which biological processes the regulated genes were involved in, gene ontology analyses were performed. Within genes upregulated in macrophages 3D cultured in low density collagen compared to 2D cultured macrophages, *cell adhesion* and *biological adhesion* were most significantly enriched (Figure 2C). Downregulated genes were especially involved in the biological processes *immune response* and *signaling* (Figure 2D).

**Figure 2.**
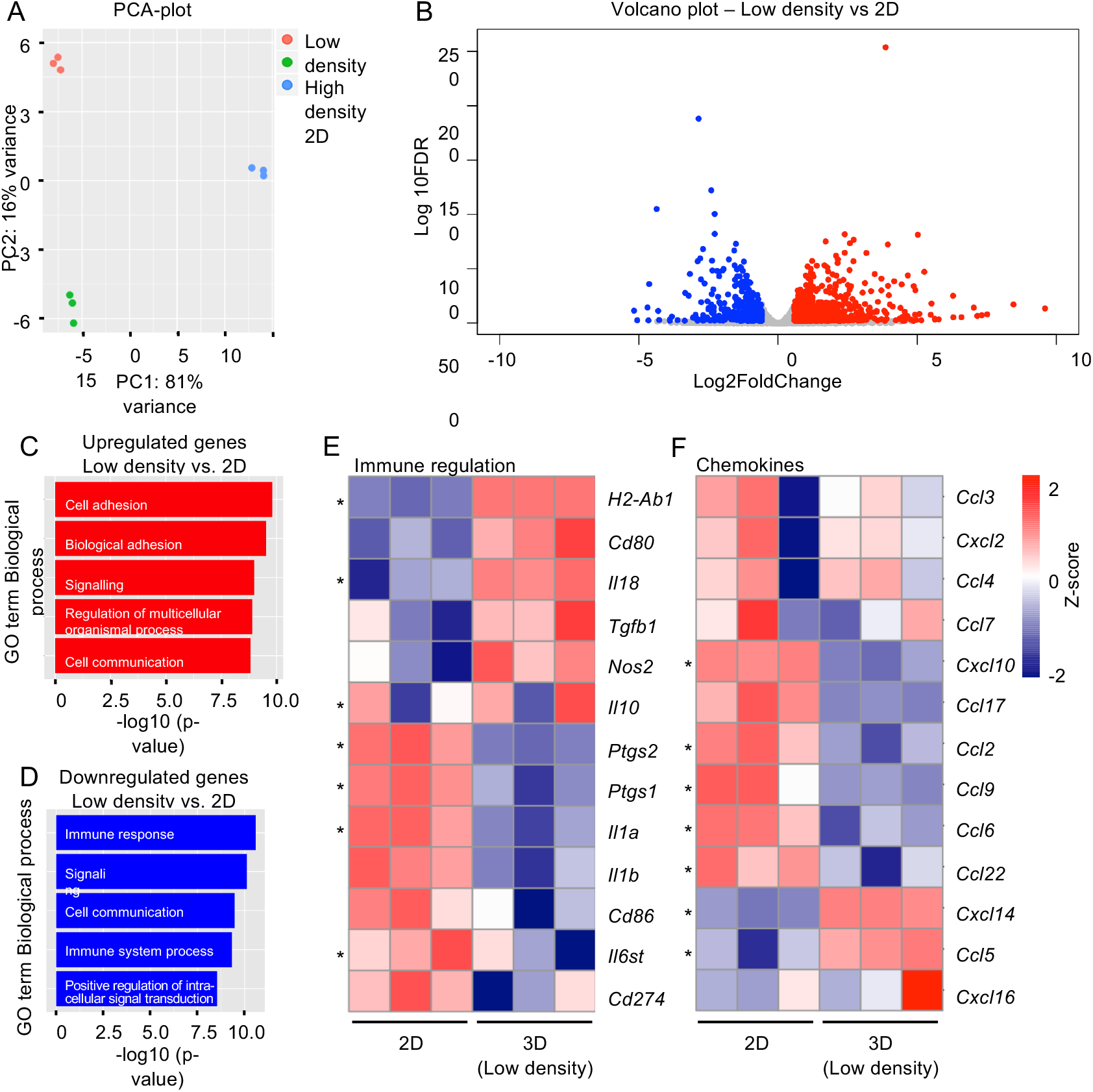
3D culture changes the gene expression profile of RAW 264.7 macrophages. RNA was extracted from RAW 264.7 macrophages cultured in collagen matrices of 1 mg/ml (Low density) or 4 mg/ml (High density) collagen or on tissue-culture treated plastic (2D) and analyzed using RNA sequencing. (A) Principal component analysis (PCA) plot of RAW 264.7 macrophages cultured in the indicated conditions. (B) Volcano plot of differentially expressed genes between RAW 264.7 macrophages grown in 3D low density collagen or on 2D tissue-culture treated plastic. Red and blue dots illustrate genes that are significantly upregulated and downregulated, respectively, in cells cultured in 3D low density collagen compared to 2D cultured cells (FDR<0.01 and Log2Fold Change > 0.585). (C and D) Gene Ontology Analysis of all differentially upregulated genes (C) or downregulated genes (D) between RAW 264.7 macrophages grown in low density collagen and on 2D tissue-culture treated plastic. (E and F) Heatmaps of the gene expression levels of a panel of selected genes involved in immune regulation (E) or encoding chemokines (F). Z-score = Normalized RPKM values. * = p-value < 0.05.

Next, the gene expression levels of a panel of immune modulatory genes and genes encoding chemokines were visualized for macrophages 3D cultured in low density collagen and 2D cultured on regular tissue-culture treated plastic surfaces (Figure 2E-F). Although several genes in both of the panels were significantly regulated, the heatmaps did not clearly indicate how the immune modulatory functions of the macrophages were affected by 3D culture compared to 2D culture (Figure 2E-F). Parallel analyses comparing samples from macrophages 3D cultured in high density collagen to 2D cultured macrophages showed that the upregulated genes were primarily involved in the biological processes *response to stimulus* and *signaling* (Figure S2B). The downregulated genes were primarily involved in the biological processes *immune response* and *immune system process* (Figure S2C). Similarly to the comparison of macrophages cultured in low density collagen to macrophages cultured in 2D, additional analysis of selected immune regulatory genes and genes encoding chemokines did not clearly show to what extent the immunosuppressive properties of the macrophages were affected (Figure S2D-E).

To investigate which transcription factor motifs could be responsible for the gene expression changes observed in 3D cultured RAW 264.7 macrophages compared to 2D cultured macrophages we used the computational method ISMARA (Integrated Motif Activity Response Analysis, https://is-mara.unibas.ch/mara/) to model transcription factor activity based on RNA sequencing data^41^. Interestingly, TEAD and SMAD transcription factor motifs were identified as upregulated in 3D cultured macrophages compared to 2D cultured macrophages (Figure S3A-B). These are known to be regulated by the mechanosensing YAP/TAZ pathway, which is activated by external mechanical cues such as extracellular matrix stiffness^42^. Consistently, several genes known to be affected by YAP/TAZ signaling were differentially regulated (Figure S3C) suggesting that YAP/TAZ signaling could be centrally engaged in the 3D culture-induced gene regulations.

### Collagen density regulates genes associated with immune regulation and chemoattraction

Next, the transcriptional differences between RAW 264.7 macrophages cultured in low-or high density collagen matrices were investigated. 385 genes were differentially regulated (FDR<0.01, Log2 fold change > 0.585) when comparing macrophages cultured in high density collagen to macrophages cultured in low density collagen. Of these, 233 genes were upregulated and 152 genes were down-regulated (Figure 3A, Figure S1C and Table 1). Gene ontology analysis showed that the upregulated genes were involved in the biological processes *defense response, inflammatory response,* and *immune response* (Figure 3B) and the downregulated genes in the biological processes *regulation of neurotransmitter levels* and *actin filament reorganization* (Figure 3C).

**Figure 3.**
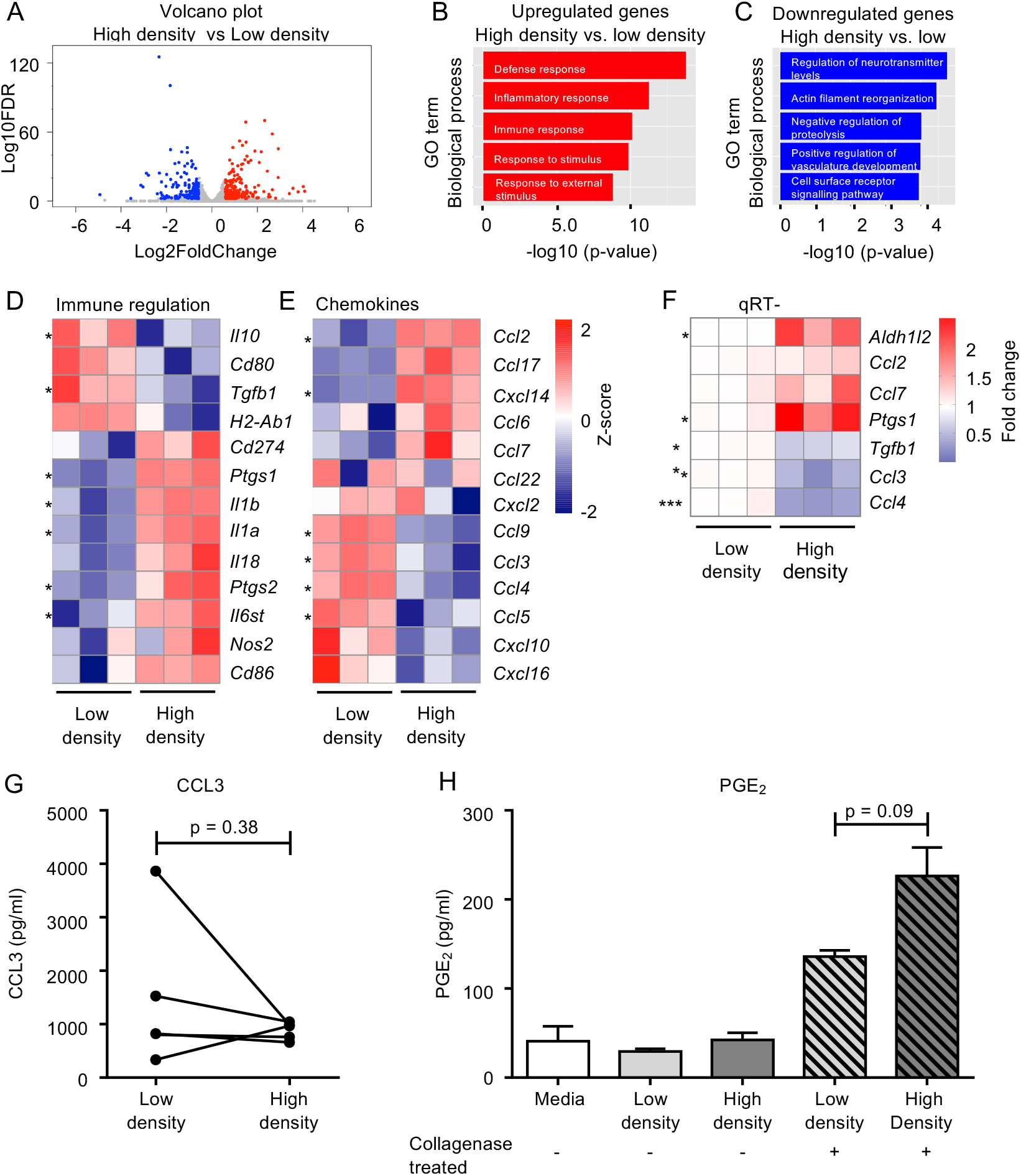
Collagen density regulates genes associated with immune regulation and genes encoding chemokines. RNA was extracted from RAW 264.7 macrophages cultured in collagen matrices of 1 mg/ml (Low density) or 4 mg/ml (High density) collagen and analyzed using RNA sequencing. (A) Volcano plot of differentially expressed genes between RAW 264.7 macrophages cultured in low- or high density collagen. Red and blue dots illustrate genes that are significantly upregulated and downregulated, respectively, in macrophages cultured in 3D high density collagen compared to macrophages cultured in low density collagen (FDR<0.01 and Log2FoldChange > 0.585. (B and C) Gene Ontology Analysis of all differentially upregulated (B) and downregulated (C) genes between RAW 264.7 macrophages cultured in low- or high density collagen matrices. (D and E) Heatmaps of the expression levels of a panel of selected genes involved in immune regulation (D) or encoding chemokines (E). Z-score = Normalized RPKM values. Asterisks indicate genes that are significantly regulated. (F) qRT-PCR analysis of RAW 264.7 macrophages cultured in low- or high density collagen. Data were obtained from three individual experiments with four samples in each condition. (G) CCL3 and (H) Prostaglandin E2 (PGE2) ELISA measurements of conditioned media and collagen-bound PGE2 (Scattered axes) from RAW 264.7 macrophages cultured in collagen matrices of low- or high density collagen. Data are means obtained from (G) five or (H) three individual experiments with four replicates per experiment. Error bars = SEM. * = p-value < 0.05. ** = p-value < 0.01. *** = p-value < 0.001.

**Table 1.**
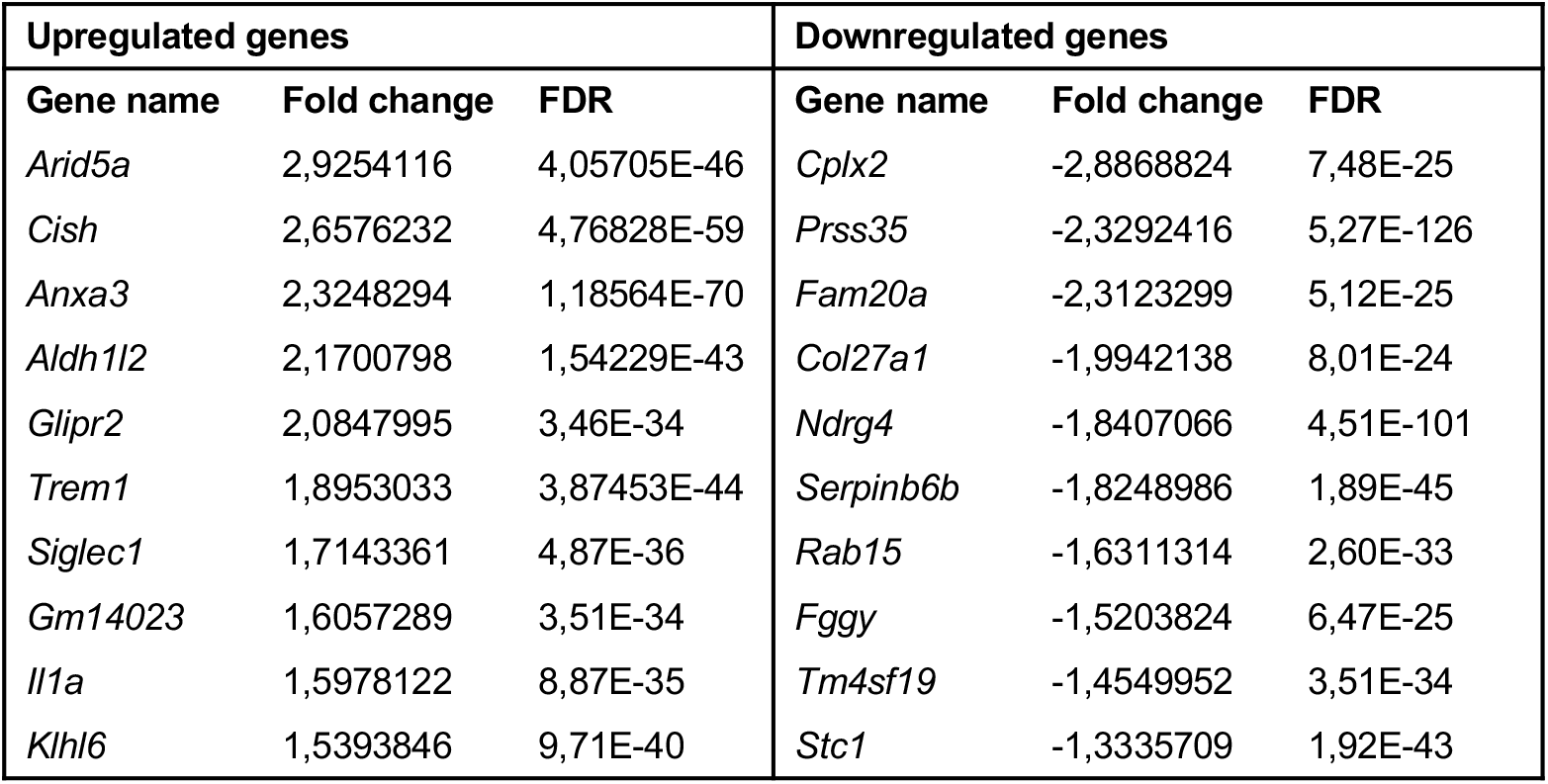
Most up- and downregulated genes between RAW 264.7 macrophages grown in high- (4 mg/ml) or low (1 mg/ml) density collagen matrices.

To further investigate if the immune modulatory activities could be affected by collagen density, we examined the gene expression levels of a panel of immune regulatory genes (Figure 3D). In line with the gene ontology analysis (Figure 3B), many of these genes were significantly regulated when comparing macrophages cultured in high density collagen to macrophages cultured in low density collagen (Figure 3D), suggesting that collagen density could affect the immune regulatory properties of macrophages. Additionally, a heatmap showing the gene expression levels of a panel of chemokines showed a clearly altered chemokine profile (Figure 3E). The transcriptional changes indicate that macrophages cultured in high density collagen would be less efficient at attracting CD8+ T cells and better at attracting T regulatory cells (Tregs) and myeloid derived suppressor cells (MDSCs), and thereby promote the generation of a more immunosuppressive microenvironment. To investigate the robustness of the RNA sequencing analysis the expression of 7 of the differentially expressed genes was tested by qRT-PCR using RNA from three independent experiments (Figure 3F). All 7 genes followed the same expression pattern but only 5 of them were significantly regulated. To investigate if the observed gene regulations also resulted in altered protein levels, conditioned media from RAW 264.7 macrophages were examined using ELISA. CCL3 is an important chemokine involved in recruitment of cytotoxic CD8+ T cells to the TME^43^. *Ptgs1* and *Ptgs2* encode the two enzymes cyclooxygenase 1 and 2 (COX-1 and −2), which are responsible for the synthesis of the prostanoid lipid, prostaglandin E2 (PGE2)^44^. In consistence with the gene expression data, we observed a trend that secretion of CCL3 was decreased and production of PGE2 was increased in RAW 264.7 macrophages cultured in high density collagen compared to low density collagen (Figure 3G-H). Interestingly, the majority of the PGE2 was not released to the conditioned media but instead bound to the collagen matrix (Figure 3H). CCL3 was not found to be associated with the collagen matrices (data not shown).

### Collagen density modulates the ability of macrophages to inhibit T cell proliferation

To investigate if the observed collagen density-induced transcriptional changes were reflected in functional changes for macrophages, we co-cultured murine splenocytes with RAW 264.7 macrophages embedded in collagen matrices of low-or high density (Figure 4A). The splenocytes and macrophages were separated by transwell inserts allowing for cell-cell communication through secreted factors. The proliferation rate of T cells was lower when co-cultured with RAW 264.7 macrophages embedded in either low- or high density collagen compared to T cells cultured alone (Figure 4B and Figure S4A). Strikingly, RAW 264.7 macrophages grown in high density collagen were capable of inhibiting T cell proliferation to a higher degree than macrophages grown in low density collagen (Figure 4B and Figure S4A). This result indicates that macrophages cultured in high density collagen matrices acquire a more immunosuppressive phenotype than macrophages cultured in low density collagen matrices. T cell proliferation were not affected when cultured above collagen matrices of low- or high density without any RAW 264.7 macrophages embedded (Figure 4B and Figure S4A). In similar co-culture assays, it was investigated if RAW 264.7 macrophages could affect the viability or phenotype of T cells. Co-culture of T cells with macrophages in different collagen densities did not affect the viability (Figure S4B) or phenotype of the T cells (Figure S4C).

**Figure 4.**
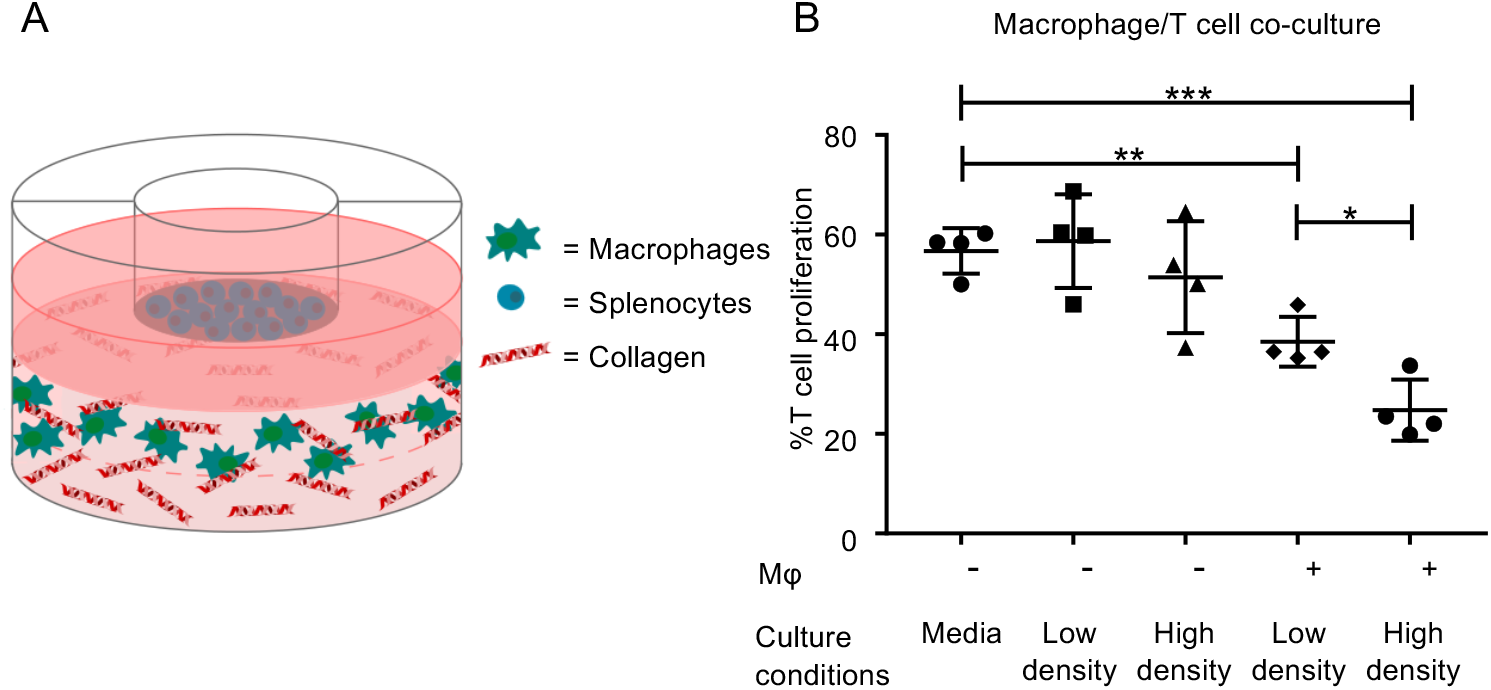
Collagen density regulates T cell proliferation through its effects on RAW 264.7 macrophages. (A) Splenocytes were isolated from the spleens of BALB/c mice. The cells were co-cultured with RAW 264.7 macrophages grown in collagen matrices of 1 mg/ml (Low density) or 4 mg/ml (High density). All samples were supplemented with 3-5 μg/ml Concanavalin A (ConA). As controls, splenocytes were cultured alone in media or above collagen matrices of low- or high collagen density without any embedded RAW 264.7 macrophages. (B) Representative example of the CD3^+^ T cell proliferation after three days of co-culture. Proliferation was measured using a BrdU based flow cytometry assay. Error bars = SD. * = p-value < 0.05. ** = p-value < 0.01. *** = p-value < 0.001.

### Collagen density alters the ability of macrophages to attract T cells

The chemokine profile of RAW 264.7 macrophages cultured in high density collagen compared to low density collagen (Figure 3E) indicated that these macrophages would be less efficient at attracting cytotoxic CD8^+^ T cells and more efficient at attracting Tregs. To investigate this possibility, T cell migration towards the conditioned media of RAW 264.7 macrophages was examined (Figure 5A). Strikingly, T cells migrated significantly less towards conditioned media from RAW 264.7 macrophages grown in high density collagen compared to conditioned media from RAW 264.7 macrophages grown in low density collagen (Figure 5B and Figure S5A). In particular, migration of CD8+ T cells towards the conditioned media from the RAW 264.7 macrophages cultured in high density collagen was clearly reduced (Figure 5C and Figure S5B) whereas CD4+ T cells were less affected (Figure 5D and Figure S5C). Interestingly, there was a trend that Tregs (CD4+/CD25+) migrated more towards the conditioned media from RAW 264.7 macrophages grown in high density collagen (Figure 5E and Figure S5D).

**Figure 5.**
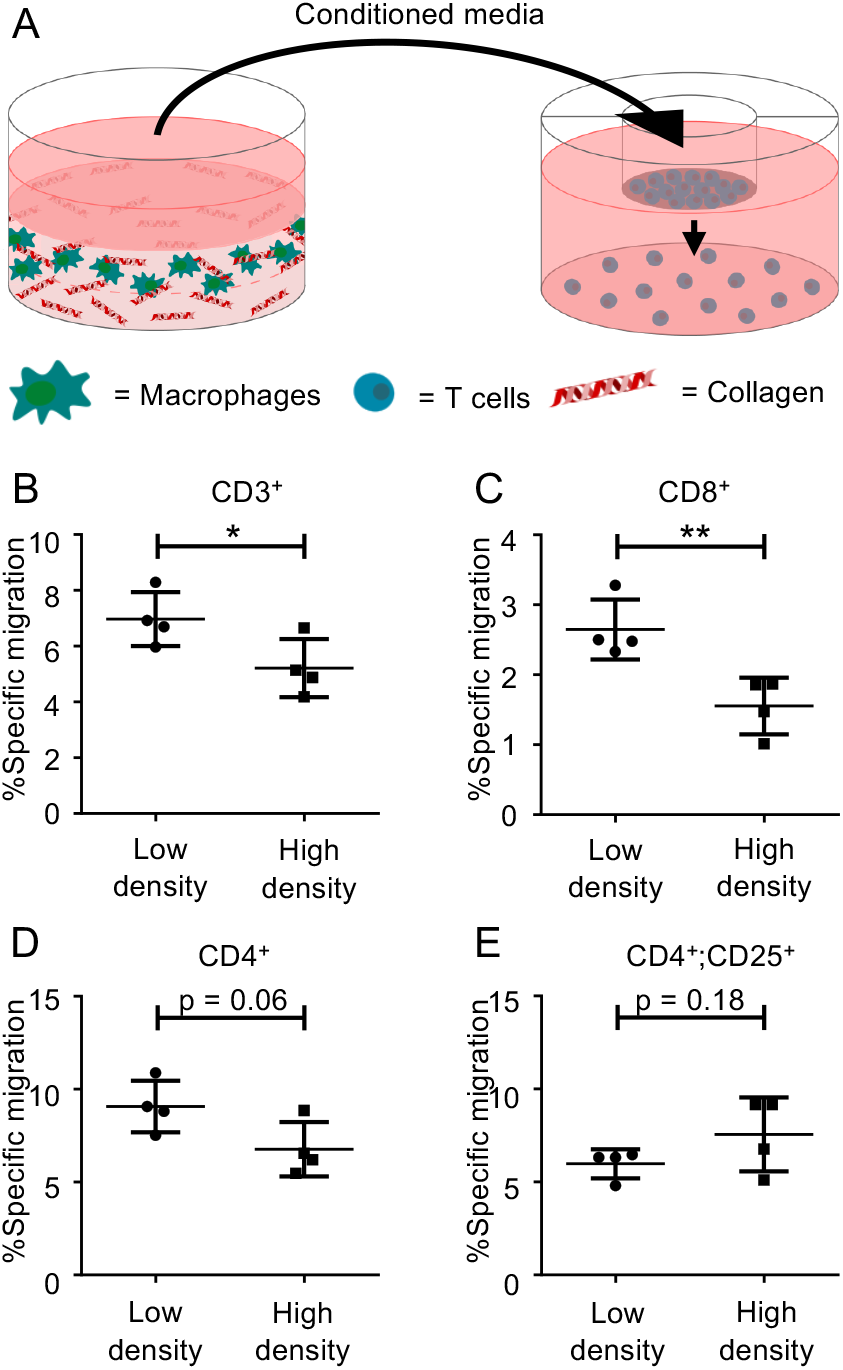
Collagen density alters the chemotactic activity of RAW 264.7 macrophages. (A) Isolated murine T cells were cultured in transwell inserts with 5 μm pore size above conditioned media from RAW 264.7 macrophages cultured in 1 mg/ml (Low density) or 4 mg/ml (High density) collagen matrices. All samples were supplemented with 3-5 μg/ml Concanavalin A (ConA). (B-E) After 26-28 hours of culture the migration of (B) CD3^+^ T cells, (C) CD3^+^;CD8^+^ T cells, (D) CD3^+^;CD4^+^CD25^+^ T cells and CD3^+^;CD4^+^CD25^+^ T cells was examined by flow cytometry. Data are representative example of the migration. Error bars = SD. * = p-value < 0.05. ** = p-value < 0.01

### Primary macrophages are affected by collagen density in a similar manner as RAW 264.7 macrophages

To investigate if the observed effects of collagen density also applied to primary macrophages, M-CSF stimulated murine BMDMs were cultured in collagen matrices of low-or high collagen density and analyzed by qRT-PCR. A panel of genes found to be significantly regulated in RAW 264.7 macrophages cultured in low-or high density collagen matrices from both RNA sequencing (Figure 3D-E and figure 6A) and qRT-PCR analysis (Figure 3F and Figure 6B) was investigated. It was found that four out of five tested genes were regulated in the same manner in BMDMs with three of these four genes reaching statistical significance (Figure 6C). To investigate if the findings also applied to TAMs, F4/80+ macrophages were isolated directly from murine 4T1 breast tumors and cultured in collagen matrices of low- or high densities. Analysis of the same panel of genes by qRT-PCR showed that the TAMs reacted to collagen density in a similar manner as RAW 264.7 macrophages and primary BMDMs (Figure 6D), although only regulation of two of the five tested genes reached statistical significance.

**Figure 6.**
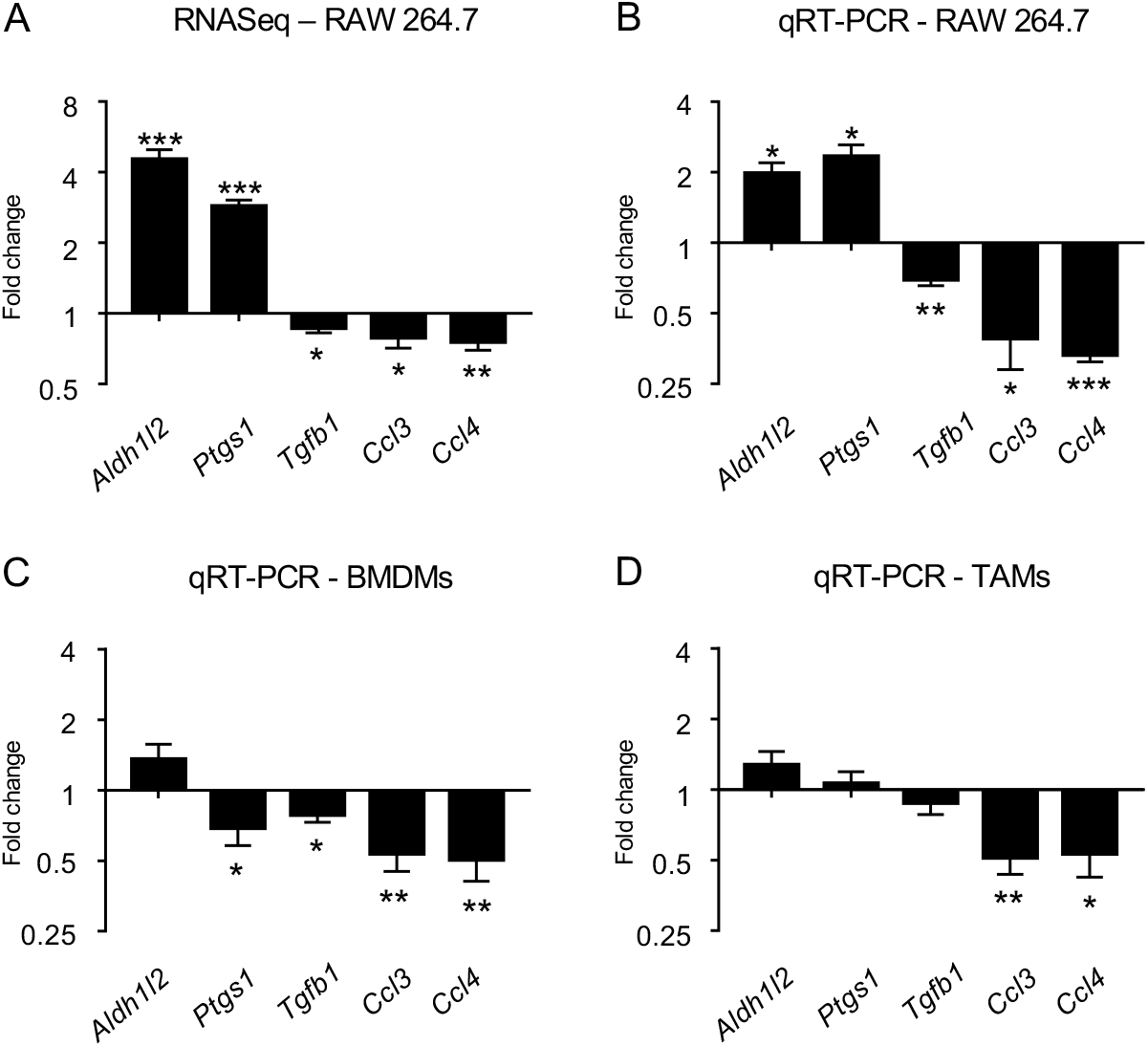
Collagen density affects the expression profile of bone-marrow derived macrophages (BMDMs) and tumor-associated macrophages (TAMs). (A) Expression of genes obtained from RNA sequencing data. RNA was extracted from RAW 264.7 macrophages cultured in collagen matrices of 1 mg/ml (Low density) or 4 mg/ml (High density) collagen. Error bars = SD (B) qRT-PCR analysis of RAW 264.7 macrophages cultured in low- or high density collagen. Data are means of three individual experiments with four samples per condition. Error bars = SEM. (C) qRT-PCR analysis of M-CSF stimulated BMDMs. M-CSF stimulated BMDMs were cultured in low- or high density collagen matrices for one day. Data are means of three individual experiments with four samples per condition. Error bars = SEM. (D) qRT-PCR analysis of F4/80+ TAMs. TAMs were isolated directly from 4T1 tumors on BALB/c mice and cultured in low- or high density collagen matrices for one day. Data are means of three individual experiments with three to four samples per condition. Error bars = SEM. * = p-value < 0.05. ** = p-value < 0.01. *** = p-value < 0.001.

## Discussion

Collagen density strongly correlates with poor prognosis in many types of cancer^8,9,11,45^. In this study, we investigated if this correlation could be due to collagen density-induced modulation of TAMs and consequently the generation of an immunosuppressive TME. Using 3D cell culture in collagen type I matrices, we examined the response of macrophages to the surrounding density of collagen.

The viability and proliferation of macrophages were not affected by 3D culture or by changes in the collagen density. However, when comparing the gene expression profile of macrophages 3D cultured in different collagen densities and in 2D, dramatic changes were nevertheless observed. The strong response is consistent with the known plasticity of macrophages and their ability to react to external stimuli^18,19^.

The comparison of macrophages cultured in high density collagen to macrophages cultured in low density collagen revealed a smaller number of differentially regulated genes than observed between macrophages cultured in 3D and in 2D. Gene ontology analysis showed that among these regulated genes the most upregulated genes were involved in the biological processes *defense response, inflammatory response,* and *immune response.* Consistently, we identified many differentially regulated genes within a selected panel of immune regulatory genes. This indicates that collagen density affects the immunomodulatory profile of macrophages. Many genes encoding chemokines were also regulated between macrophages cultured in low- or high density collagen matrices. The observed changes suggested a reduced recruitment of cytotoxic CD8+ T cells (downregulation of *Ccl3, Ccl4, Ccl5, Cxcl10* and *Cxcl16)* and an increased recruitment of Tregs, MDSCs, macrophages and monocytes (upregulation of *Ccl2, Ccl6, Ccl7, Ccl17* and Cxcl14)^43–46,47^. By investigating a selected panel of the regulated genes, we demonstrated that primary BMDMs and TAMs respond in a similar manner to the surrounding collagen density. This indicates that the effect of collagen density applies to various subsets of macrophages, and thus substantiates the importance of the ECM structure for macrophage phenotype and function.

To investigate how these transcriptional changes affected the functions of the macrophages, migration assays to examine the chemoattraction of T cells were conducted. As hypothesized based on the chemokine profile, macrophages cultured in high density collagen were less efficient at attracting cytotoxic CD8+ T cells. Stromal collagen has been suggested to prevent the migration of cytotoxic T cells into the tumor islets^48,49^. Based on our observed changes of the chemokine profile of macrophages, we speculate that a high collagen density could also limit T cell influx by reducing the macrophages ability to attract cytotoxic T cells. Interestingly, co-culture assays also showed that macrophages cultured in a high density collagen matrix become capable of inhibiting T cell proliferation to a greater extent than macrophages cultured in low density collagen. This is the first demonstration that the collagen density can directly modulate the immunosuppressive activity of macrophages. The identified mechanism could be important for the generation of an immunosuppressive TME.

The reason why macrophages cultured in high density collagen inhibit T cell proliferation more than macrophages cultured in low density collagen still needs to be investigated. ELISA analysis showed that macrophages generate more PGE2 when cultured in high density collagen. PGE2 is known to affect T cell activity and phenotype^50–52^ and could be one of the factors involved in the observed suppression of T cell proliferation.

Recently, we have demonstrated that a high collagen density can also directly downregulate the cytotoxic activity of T cells and instead make them more regulatory^15^. Together with the findings of this study, these observations suggest that the collagen density within tumors could be centrally engaged in the formation of an immunosuppressive TME and thus be one of the mechanisms limiting the efficacy of immunotherapy. This knowledge could potentially be used for identifying non-responsive groups of patients and to tailor therapeutic interventions. Additionally, the results indicate that the prognostic value of collagen density could be related to its immune regulatory effects on immune cells such as macrophages.

Exactly how collagen density exerts its effect on macrophages still needs to be investigated. One possibility is that macrophages cultured in high density collagen are affected through the increased stiffness of the matrix. Collagen matrices of 4 mg/ml are stiffer than collagen matrices of 1 mg/ml^16,40^ and studies have indicated that macrophages cultured on stiff substrates acquire a more anti-inflammatory phenotype^53–55^. Another possibility is that the increased number of binding sites for collagen receptors found in the high density collagen matrices could potentiate collagen-induced signaling. In this regard, it could be interesting to investigate the role of different collagen receptors, including β1-integrins, discoidin domain receptor tyrosine kinase 1 (DDR1), DDR2, osteoclast-associated receptor (OSCAR), leukocyte-associated immunoglobulin-like receptor 1 (LAIR-1), and mannose receptor (MR)^56,57^. It also remains to be elucidated which intracellular signaling pathways are responsible for the collagen-induced modulation of macrophages. YAP/TAZ signaling was in this study identified as one of the potential pathways. Gaining a better understanding of the molecular mechanisms underlying the response of macrophages to the surrounding collagen density could reveal targets for new cancer therapy. Such therapy could prevent the generation of immunosuppressive macrophages and thereby enhance the efficacy of existing immunotherapy.

Increased mammographic density is a risk factor for the development of cancer and has been associated with increased collagen density^58^. Additionally, studies in mice^17,59^ and humans^60^ have demonstrated that increased collagen density can promote tumor progression. In this study, we have identified collagen type I as a novel regulator of macrophage function, indicating that the effect of collagen density on tumor progression could be caused by its impact on macrophage function. A study by Pinto and colleagues^61^ investigated the effects of tumor ECM on macrophages by growing the cells on decellularized matrices from normal tissue sections and colorectal cancer tissues. Macrophages grown on tumor matrices acquired a more anti-inflammatory phenotype. These findings are in line with the results obtained in this current study. However, the ECM is a complex network composed of various macromolecules including other types of collagen as well as many non-collagen ECM proteins. Many of these components of the ECM could potentially also influence the immunosuppressive activity of macrophages. A recent study demonstrated that macrophages co-cultured with versican deficient mesothelioma cells acquired a more pro-inflammatory phenotype with increased phagocytic activity^62^. Versican is an ECM component that is upregulated in cancers^63–65^, and it has been suggested to have immunosuppressive functions^66,67^. Other studies have identified the ECM component hyaluronan as being capable of stimulating TAMs to acquire an immunosuppressive M2-like phenotype^68,69^. These findings specify the importance of also investigating the effect of other non-collagen ECM components on the function of macrophages.

In conclusion, we have identified collagen density as an important regulator of the immunosuppressive phenotype of TAMs. This mechanism could be critical for the ability of cancers to evade immune destruction and could constitute a limiting factor for the efficacy of immunotherapy.

## Acknowledgments

This study was supported by the Danish Cancer Society (D.H.M.), Knæk Cancer (D.H.M.), Novo Nordisk Foundation (D.H.M.), Dagmar Marshalls Foundation (D.E.K., D.H.M., A.M.H.L), Dansk Kræftforskningsfond (D.E.K.), Einar Willumsen Foundation (D.E.K.), Jens og Maren Thestrups legat til kræftforskning (A.M.H.L.), Agnes og Poul Friis’ Fond (A.M.H.L.), and Herlevs Forskningsfond (A.M.H.L.). We thank Dr. Janine T. Erler for providing BALB/c mice.

## Author contributions

Conceptualization, D.H.M. and A.M.H.L.; Methodology, D.H.M., A.M.H.L., D.E.K., L.G., and O.V.; Investigation, D.H.M., A.M.H.L., D.E.K., A.K., M.S.S., M.L.T., M.C., A.Z.J.; L.G.; Writing – Original Draft, D.H.M. and A.M.H.L.; Writing – Review & Editing, D.H.M., A.M.H.L., D.E.K., A.K., M.S.S., M.L.T., M.C., A.Z.J., L.G., and O.V.

