## Supplementary for "Collagen Density Modulates the Immunosuppressive Functions of Tumor-Associated Macrophages"

**Supplementary table 2.** Primer sequences used for qRT-PCR.

| **Gene name** | **Forward strand** | **Reverse strand** | **Primer efficiency** |
|---|---|---|---|
| *Actb* | CACTGTCGAGTCGCGTCC | TCATCCATGGCGAACTGGTG | 97.4% |
| *Aldh1l2* | TCCATCATCTACCACCCCTC | TCCCATGATTAGTGTCCAGTTG | 101.22% |
| *Tgfb1* | AGCCCTGTATTCCGTCTCCT | CTGCTGACCCCCACTGATAC | 110.28% |
| *Ptgs1* | CTGGAGTTGCACCCGAGG | GCGAGAGACTCCTTCGACTC | 95.71% |
| *Ccl3* | TACAGCCGGAAGATTCCACG | TCAGGAAAATGACACCTGGCT | 108.5% |
| *Ccl2* | GCTGTAGTTTTTGTCACCAAG | TATGTCTGGACCCATTCCTT | 97% |
| *Ccl4* | AGCTGTGGTATTCCTGACCA | ACTCCAAGTCACTCATGTACT | 86.34% |
| *Ccl7* | CACATGCTGCTATGTCAAGA | CTTGAAGATAACAGCTTCCCAG | 97.74% |

**Supplementary table 3.** Most up- and downregulated genes between RAW 264.7 macrophages grown in 3D low density (1 mg/ml) collagen matrices or on 2D tissue-culture treated plastic.

| **Upregulated genes** | | | **Downregulated genes** | | |
|---|---|---|---|---|---|
| **Gene name** | **Fold change** | **FDR** | **Gene name** | **Fold change** | **FDR** |
| *Cplx2* | 9,5924134 | 3,59E-14 | *Arid5a* | -4,3597695 | 1,00E-105 |
| *Prss35* | 8,4595737 | 9,31E-18 | *Cish* | -2,8873012 | 1,72E-57 |
| *Fam20a* | 4,8772656 | 9,42E-35 | *Lpl* | -2,3963320 | 6,19E-123 |
| *Siglec1* | 4,4406316 | 2,72E-36 | *Aldh1l2* | -2,3570131 | 3,71E-49 |
| *Col27a1* | 4,1713975 | 3,15E-39 | *Icosl* | -2,2741569 | 3,83E-101 |
| *Rab15* | 3,9468267 | 4,43E-73 | *Anxa3* | -2,1923171 | 8,54E-49 |
| *Paqr5* | 3,5432448 | 8,22E-12 | *Gm14023* | -1,5298722 | 1,00E-23 |
| *Scn11a* | 2,7191268 | 1,78E-77 | *Il1a* | -1,5262392 | 2,96E-24 |
| *Stc1* | 2,6049839 | 2,26E-13 | *Ctsc* | -1,4552543 | 9,09E-44 |
| *Fggy* | 2,5109535 | 1,13E-35 | *Fat1* | -1,3328074 | 2,58E-20 |

**Supplementary table 4.** Most up- and downregulated genes between RAW 264.7 macrophages grown in 3D high density (4 mg/ml) collagen matrices or on 2D tissue-culture treated plastic.

| **Upregulated genes** | | | **Downregulated genes** | | |
|---|---|---|---|---|---|
| **Gene name** | **Fold change** | **FDR** | **Gene name** | **Fold change** | **FDR** |
| *Siglec1* | 6,1460430 | 3,42E-84 | *Pde6b* | -2,9367260 | 8,89E-62 |
| *Prss35* | 6,1388278 | 4,91E-09 | *Hpgds* | -2,1347715 | 2,75E-84 |
| *Nt5e* | 4,6595411 | 1,95E-38 | *Gadd45b* | -2,1275461 | 3,04E-82 |
| *Glipr2* | 3,6025141 | 1,64E-55 | *Pgm5* | -2,1249075 | 9,00E-31 |
| *Axl* | 3,0743984 | 1,03E-154 | *Lpl* | -1,5960209 | 3,47E-57 |
| *Paqr5* | 2,4597318 | 5,03E-06 | *Arid5a* | -1,4383000 | 1,60E-24 |
| *Rab15* | 2,3249462 | 2,19E-18 | *Serpinb6b* | -1,4170062 | 7,53E-22 |
| *Sema4c* | 1,8058068 | 2,79E-43 | *Igf2bp2* | -1,3198774 | 1,07E-42 |
| *Scn11a* | 1,5194497 | 6,54E-18 | *Acy1* | -1,2933965 | 2,29E-17 |
| *Scd1* | 1,4303213 | 2,54E-63 | *Gla* | -1,1487874 | 7,59E-24 |

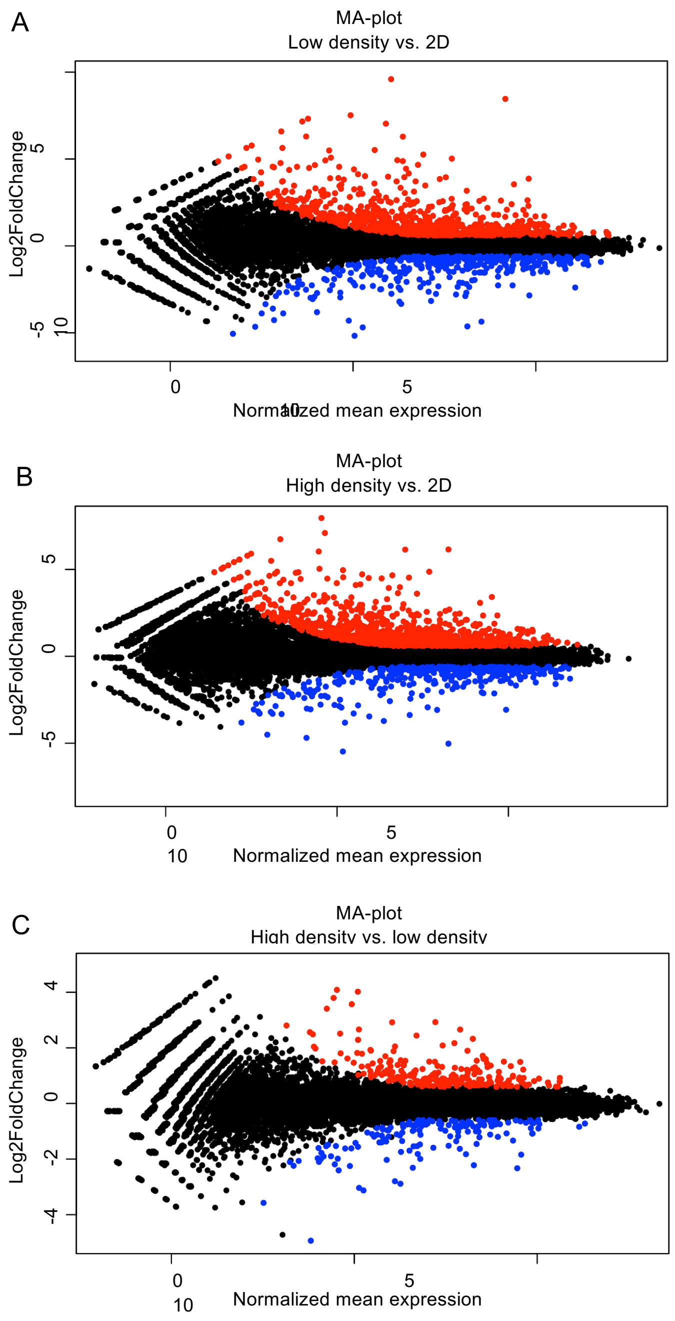


**Supplementary figure 1.** (A) MA-plot of differentially regulated genes between RAW 264.7 macrophages cultured in 1 mg/ml (Low density) collagen matrices or on tissue-culture treated 2D plastic (2D). (B) MA-plot of differentially regulated genes between RAW 264.7 macrophages cultured in 4 mg/ml (High density) collagen matrices or on tissue-culture treated 2D plastic. (C) MA-plot of differentially regulated genes between RAW 264.7 macrophages cultured in low- or high density collagen matrices. Red and blue dots illustrate genes that are significantly up- or downregulated, respectively (FDR<0.01 and Log2Fold Change > 0.585).

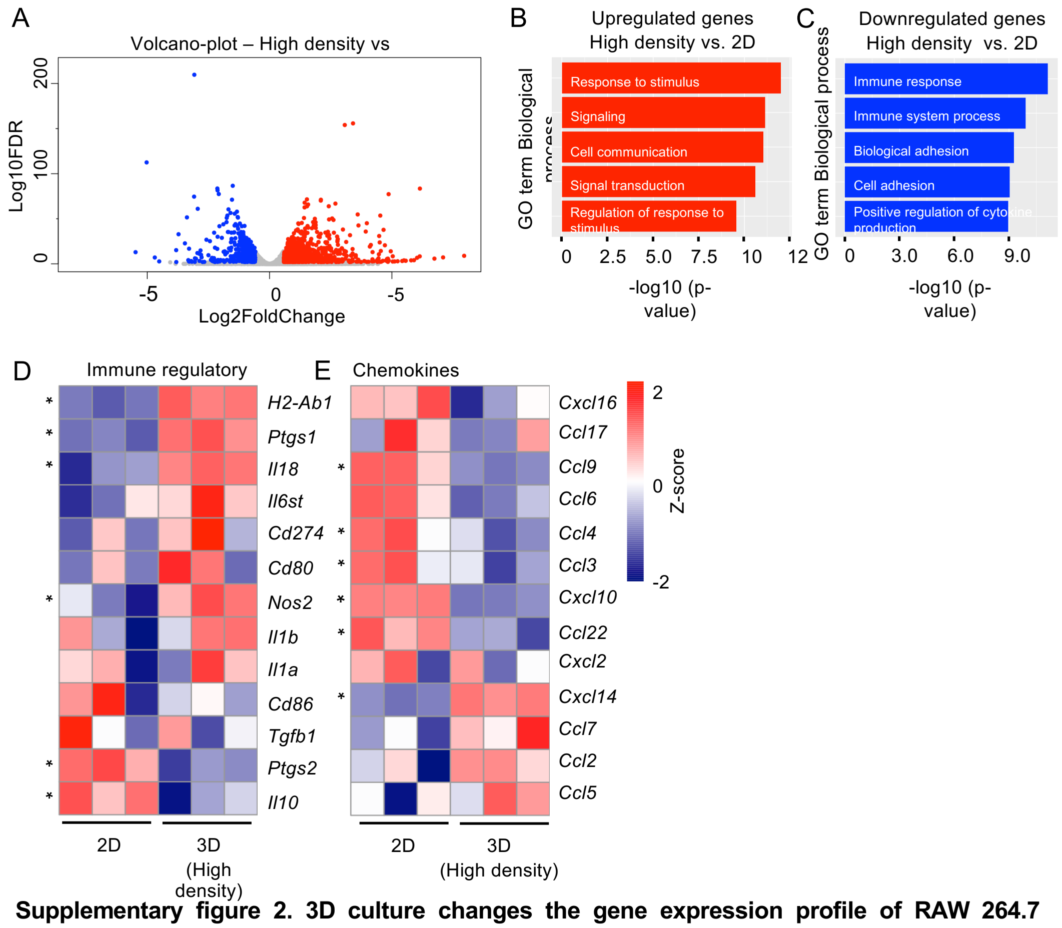


**Supplementary figure 2. 3D culture changes the gene expression profile of RAW 264.7 macrophages.** RNA was extracted from RAW 264.7 cells cultured in collagen matrices of 1 mg/ml (Low density) or 4 mg/ml (High density) collagen or on tissue-culture treated plastic (2D) and analyzed using RNA sequencing. (A) Volcano plot of differentially expressed genes between RAW 264.7 macrophages grown in 3D high density collagen or on 2D tissue-culture treated plastic. Red and blue dots illustrate genes that are significantly up- or downregulated, respectively (FDR<0.01 and Log2Fold Change > 0.585). (B and C) Gene Ontology Analysis of all differentially upregulated genes (B) or downregulated genes (C) between RAW 264.7 macrophages grown in high density collagen or on 2D tissue-culture treated plastic. (D and E) Heatmaps of the gene expression levels of a panel of selected genes involved in immune regulation (D) or encoding chemokines (E). Z-score = Normalized RPKM values. * = p-value < 0.05.

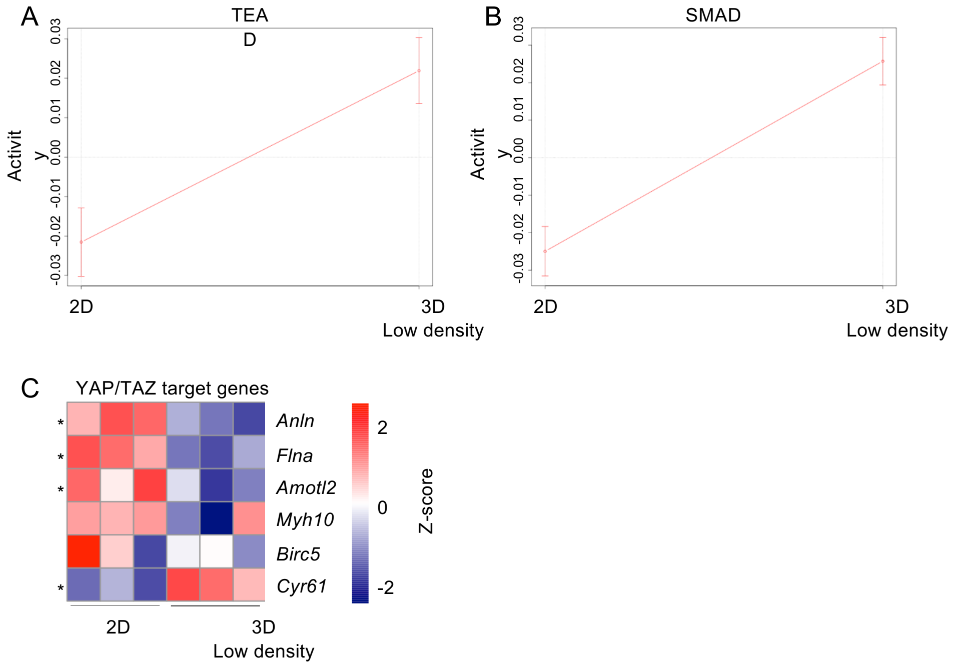


**Supplementary figure 3. YAP/TAZ signaling could be centrally engaged in the 3D culture-induced gene regulations.** (A-C) RNA was extracted from RAW 264.7 macrophages cultured in collagen matrices of 1 mg/ml (Low density) or 4 mg/ml (High density) collagen or on tissue-culture treated plastic (2D) and analyzed using RNA sequencing. (A-B) Selected top ranked transcription factors found using from ISMARA analysis of significantly regulated genes between RAW 264.7 macrophages grown in 3D low density collagen and 2D tissue-culture treated plastic. (C) Heatmaps of the expression levels of a panel of selected YAP/TAZ target genes. Z-score = Normalized RPKM values. * = p-value < 0.05.

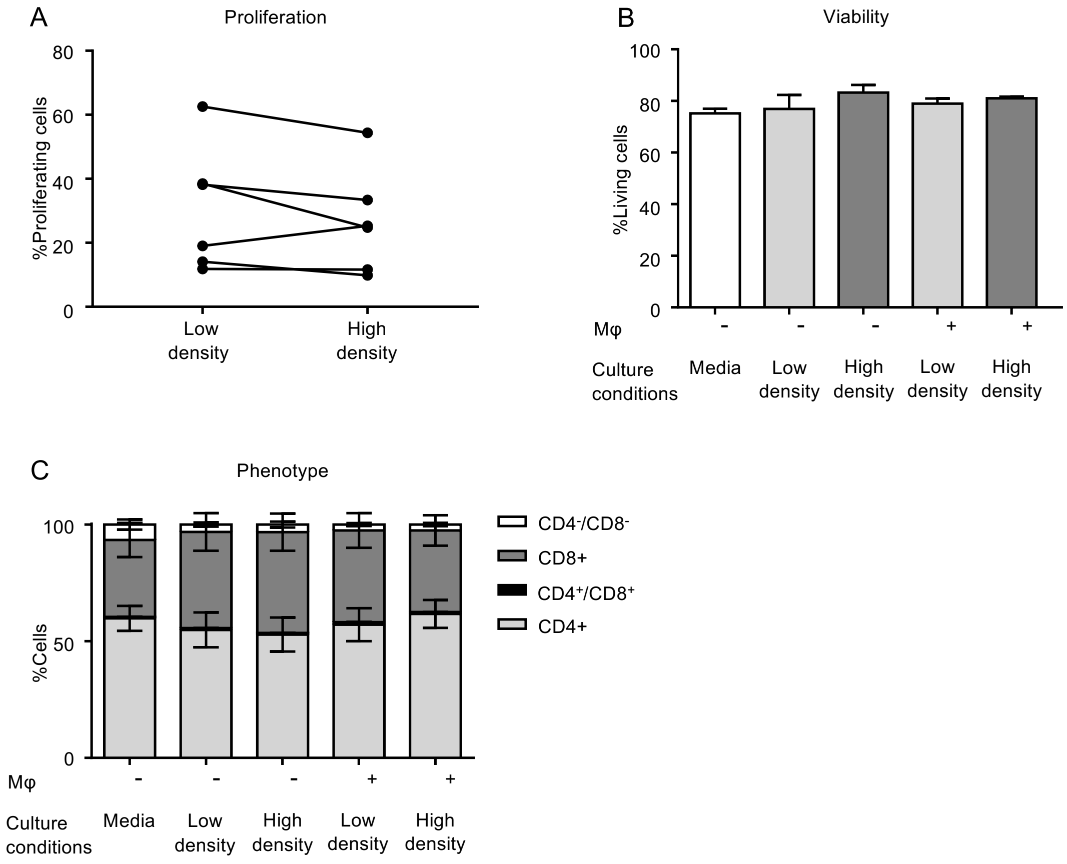


**Supplementary figure 4. Collagen density regulates T cell proliferation but not T cell viability and phenotype through its effects on RAW 264.7 macrophages.** (A-C) Splenocytes were isolated from the spleens of BALB/c mice. The cells were co-cultured with RAW 264.7 macrophages grown in collagen matrices of 1 mg/ml (Low density) or 4 mg/ml (High density). All samples were supplemented with 3-5 µg/ml Concanavalin A (ConA). As controls, splenocytes were cultured alone in media or above collagen matrices of low- or high collagen density without any embedded RAW 264.7 macrophages. After three days of co-culture the proliferation (A), viability (B) and phenotype (C) of $CD3^+$ T cells were evaluated using a BrdU based flow cytometry assay. Data are means obtained from (A) six and (B-C) three experiments with three-four samples per condition. Error bars = SEM.

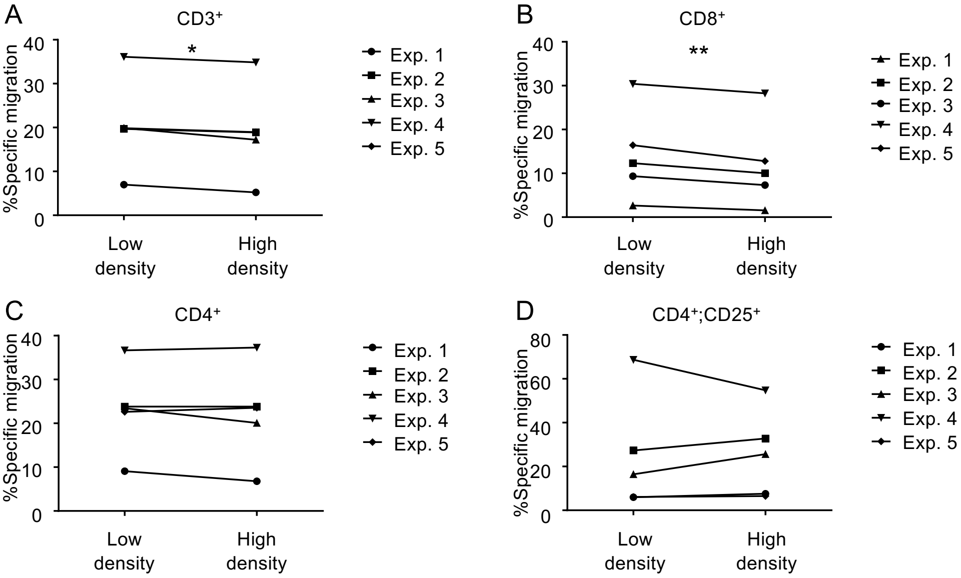


**Supplementary figure 5.** (A-D) Isolated murine T cells were cultured in transwell inserts with 5 µm pore size above conditioned media from RAW 264.7 macrophages cultured in 1 mg/ml (Low density) or 4 mg/ml (High density) collagen matrices. All samples were supplemented with 3-5 µg/ml Concanavalin A (ConA). After 26-28 hours of culture the migration of (A) $CD3^+$ T cells, (B) $CD3^+$;$CD8^+$ T cells, (C) $CD3^+$;$CD4^+CD25^-$ T cells and (D) $CD3^+$;$CD4^+CD25^+$ T cells was examined by flow cytometry. Data are means from five individual experiments with four samples per condition. * = p-value < 0.05. ** = p-value < 0.01
